# Palaeogenomics suggest domesticated camelid herding and wild camelid hunting in early pastoralist societies in the Atacama Desert

**DOI:** 10.1101/2024.10.09.617380

**Authors:** Conor. O’Hare, Paloma Diaz-Maroto, Isabel Cartajena, Michael. V Westbury

## Abstract

The domestication of camelids played a significant role in the development of Andean societies. Although South American camelids can be osteometrically classified as large (putatively *Lama*), or small (putatively *Vicugna*), it is difficult to differentiate between wild or domesticated. Here we utilise palaeogenomic data to reveal the sex and ancestry of camelids from archaeological sites from the Tulán ravine, a key area for understanding the transition from hunter-gatherer to pastoral societies in the South-Central Andes. We inferred the ancestry of 49 individuals with genome-wide coverages >0.001x and found evidence for both *Vicugna* and *Lama*. Investigations into 26 individuals of >0.01x showed that all individuals, except one male llama, represented ancestry not found in modern individuals. Similar male-to-female sex ratios suggest hunting rather than herding. Moreover, while intergeneric admixture is widespread in modern domesticated individuals, we find limited evidence of this in our dataset. Overall, our findings suggest that lost early domesticate lineages and/or wild camelids, not the direct ancestors of contemporary domesticates, were the primary source of camelids in both domestic and ritual contexts in the Tulán ravine during the Early Formative period (3,360–2,370 cal. yr BP).

## Introduction

One of the most significant cultural developments of humanity was the domestication of plants and animals ^1^. Domestication refers to the process by which humans selectively breed wild animals and plants for specific traits that serve human needs, leading to genetic, morphological, and behavioral changes over generations ^2^. As domesticated animals became central to human societies, pastoralism emerged as a subsistence strategy, where humans raise and manage these animals for resources such as food, fiber, and transport. However, despite its remarkable significance, domestication processes (e.g. wild ancestors, timing, location, and underlying genetic mechanisms) remain unsolved for many species ^3^. This is especially true for the only large herd animals domesticated in the Americas, South American camelids ^1^. The domestication of South American camelids played a significant role in the development of Andean societies. Their lone domesticated status and ability to occupy broad domesticate roles (transport, fibre, food) exemplifies the pivotal role these animals played in shaping ancient South American societies. Camelid fibres and meat served both practical and ritual purposes, while pack llama (*Lama glama*) acted to connect distant communities ^4–6^. Even today, South American camelids remain the most important resource for many communities ^7^.

South American camelids comprise four extant species; two domesticated (llama [*Lama glama*], alpaca [*Vicugna pacos*]), and two wild (guanaco [*L*.*guanaco*], vicuña [*V. vicugna*]). Within the wild species there are two subspecies each: *L*.*g*.*cacsilensis, L*.*g*.*guanicoe, V*.*v*.*mensalis*, and *V*.*v*.*vicugna*. While multiple regions show evidence of early camelid management, such as hunting, herd protection, and the early stages of captive care, the most likely centers of domestication, where the controlled breeding of camelids began, are currently considered to be the Central Andes, Southern Bolivia, and the Circumpuna region spanning Northern Chile and NW Argentina^8–14^. However, the timing and number of domestication centres remain debated ^15^. Archaeological evidence suggests South American camelid domestication was a complex process and not synchronous across the Andes that began with hunting, herd protection, and finally captivity/selective breeding ^16^. However, although South American camelids can be osteometrically classified as large (putatively *Lama*), or small (putatively *Vicugna*), it is difficult to designate an individual as wild or domesticated ^17,18^. This is primarily due to the significant overlap in size between wild and domesticated forms, as well as the lack of clear, consistent morphological differences between them. These factors complicate efforts to definitively classify individuals as wild or domesticated, limiting insights into the presence of early domesticated individuals.

Genetics is a useful tool that can complement the archaeological record to provide further insights into domestication ^19^. However, despite attempts to address theories of South American camelid domestication, the Spanish conquest of South America beginning ∼1500 A.D and uncontrolled post-conquest intergeneric hybridisation between domesticated species makes it difficult to identify genetic affinities prior to ∼1500 A.D from contemporary individuals ^20^. This intergeneric hybridization, which is thought to have become widespread following the Spanish arrival, has significantly shaped the genetic makeup of modern domesticated llama and alpaca populations ^20^. As a result, modern populations are highly admixed, complicating efforts to trace pure genetic lineages that could shed light on the early domestication process. Ancient DNA (aDNA) from pre-conquest individuals is therefore a useful approach to reconstructing the uncertain past of South American camelid domestication and husbandry. However, although previous studies have utilised pre-conquest individuals, these have been limited to mitochondrial DNA ^17,21–23^. Mitogenomes are maternally inherited, only represent a single locus, and are confounded by hybridisation and stochastic events ^24^. This can be especially problematic in the case of South American camelids as modern individuals are known to contain high levels of hybridisation ^20^, limiting the power of mitochondrial reference panels for taxonomic identifications.

Here we utilise palaeogenomic data to investigate the use of camelids by ancient South American communities. We analyse the sex and ancestry from pre-conquest South American camelids recovered from archaeological sites from the Tulán ravine (Tulán-52, Tulán-54, Tulán-85, and Tulán-94) located in the eastern border of the Salar de Atacama that descends into Northern Chile (Fig 1). The Tulán ravine is a key area for the transition from hunter-gatherer societies, which are characterised by subsistence through hunting, fishing, and foraging of wild plants, to initial pastoral societies, where domesticated animals are herded for food, labour, and other resources, in the South-Central Andes ^25^. The sites studied here belong to the Late Archaic period (Tulán-52; 5290-4840/4430-4090 cal. yBP) and to the Early Formative period (Tulán-54, Tulán-85, and Tulán-94; 3,360–2,370 cal. yr BP) when domesticated camelids are thought to have been incorporated into the economic activities of humans of this area. Specifically, archaeological remains from ritual (sites associated with ceremonial or religious practices) and domestic (sites used for daily living and subsistence activities) contexts provide evidence of a significant cultural and economic transformation from in the highlands of the Southern Andes. ^26^.

**Figure 1:**
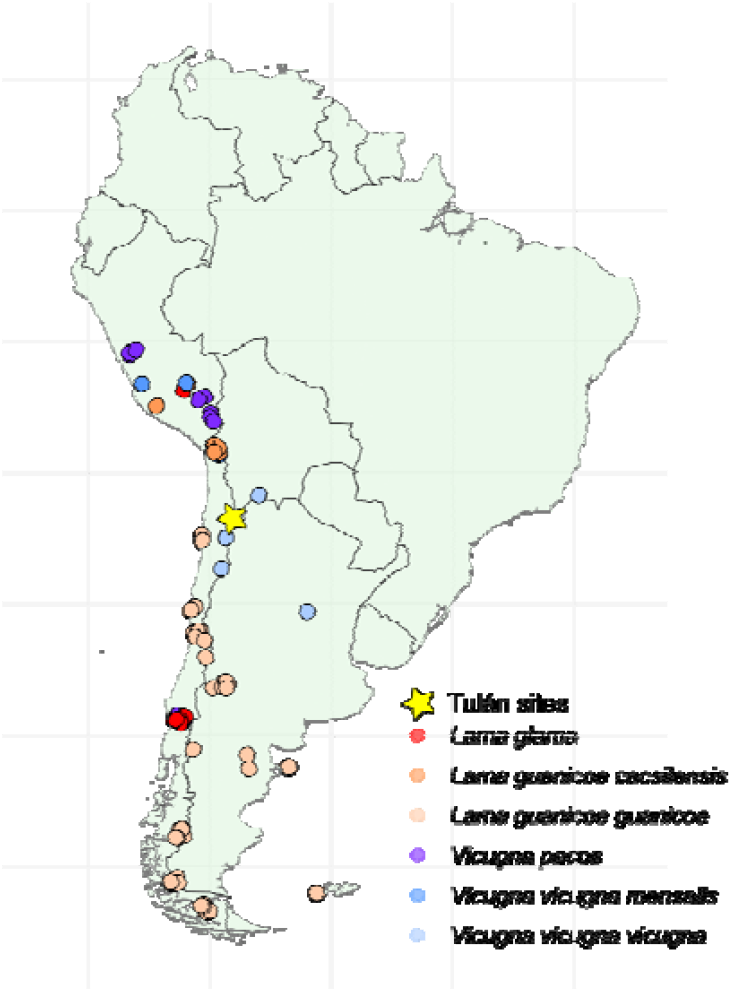
Map indicating the location of the Tulán archaeological sites and the sampling location of the previously published modern South American camelid individuals used in this study.

## Results

### Mapping results

Our study utilised genomic data from a total of 75 ancient camelid individuals, sourced from four archaeological sites: Tulán-52, Tulán-54, Tulán-85, and Tulán-94. These individuals had previously undergone mitochondrial and morphological analysis (Supplementary Tables S1 and S2) but have yet to be analysed from a nuclear genomics perspective. After mapping to the alpaca reference genome, genome-wide coverages ranged between ∼0 and 0.28x or 4,500 and 587 million mapped base pairs (Supplementary table S2). The ancient samples showed typical ancient DNA damage patterns with elevated levels of A-G and C-T transitions towards the read ends (Supplementary Fig S1).

To build a genomic reference panel, we also incorporated previously published genomic data from 80 modern individuals representing all extant *Lama* and *Vicugna* subspecies, which served as a comparative basis for the ancient samples. After mapping to the alpaca reference genome, all modern individuals had genome-wide coverages ranging from 8.34x to 41.11x (Supplementary table S1).

### Principal Component Analysis

To evaluate the broad relationships between the modern and ancient camelids included in this study, we performed principle component analyses (PCAs) using a pseudohaploid base call with ANGSD. When analysing the entire dataset (*Lama sp*. and *Vicugna sp*.), the modern individuals fell into five distinct clusters, in most cases pertaining to their subspecies classification (Fig 2). However, we found *L*.*g*.*cacsilensis* and llamas clustering together, apart from two *L*.*g*.*cacsilensis* which clustered with *L*.*g*.*guanicoe* (Cacsilensis1 and Cacsilensis2). These two individuals are from outside of the known range of *L*.*g*.*cacsilensis* and were therefore likely misidentified ^20^.

**Figure 2:**
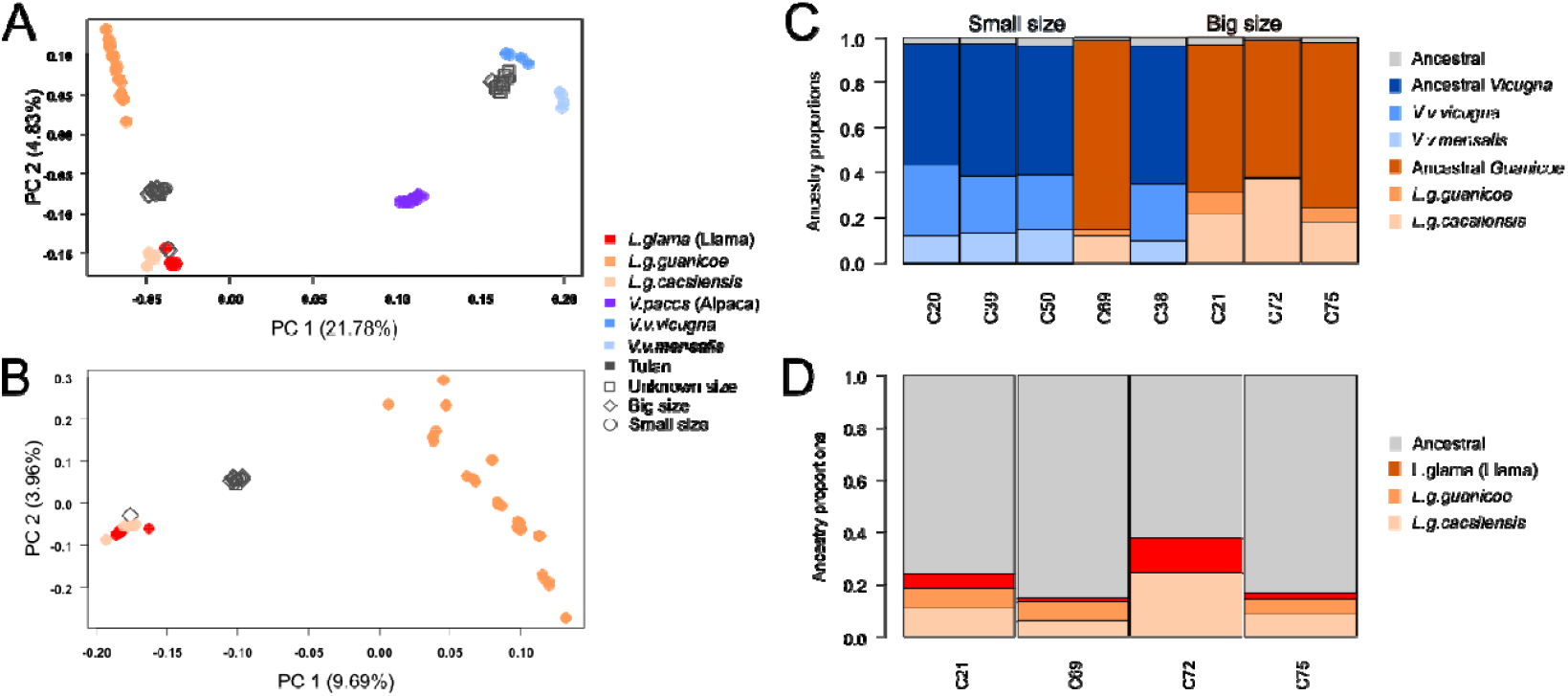
Relationships between modern and ancient individuals. A) PCA performed on all modern individuals and ancient individuals >0.01x using 5,163,744 transversion sites. B) PCA performed on modern *Lama* individuals and ancient *Lama* individuals > 0.01x using 3,668,867 transversion sites. C) Ancestry states of eight ancient individuals. Ancestry states are derived from a single individual; *L*.*g*.*cacsilensis* - Cacsilensis4, *L*.*g*.*guanicoe* - Guanicoe1, *V*.*v*.*mensalis* - Mensalis4, and *V*.*v*.*vicugna*. Ancestral is defined as having combined ancestry between both *Lama* and *Vicugna* (C) or mixed ancestry between *Lama* (sub)species (D).

When considering the ancient samples >0.01x, we see three groupings; One closest to *V*.*v*.*vicugna* and mostly containing osteometrically small size individuals, one mostly containing osteometrically large size individuals somewhat in the middle of the two modern *Lama sp*. clusters, and a single ancient individual (C72) within the llama/*L*.*g*.*cacsilensis* cluster (Fig 2A). When reducing the dataset to only include *Lama sp*. individuals, we again see C72 within the llama/*L*.*g*.*cacsilensis* cluster, with the remaining individuals still forming their own cluster, but sitting more closely to the llama/*L*.*g*.*cacsilensis* cluster (Fig 2B). Results are similar when considering ancient individuals >0.001x; the cluster closest to *V*.*v*.*vicugna* remains but the structuring within *Lama sp*. becomes less clear (Supplementary Fig S2). Individuals where the osteometric size did not match with their PCA placement included a single big individual (C38), and two small individuals (C69 and C70).

### Phylogenetic Analysis

To further investigate the broad relationships between the modern and ancient camelids included in this study, we computed a neighbour joining tree. Our neighbour joining tree showed similar results as the PCA (Supplementary Fig S3). There is a split in the ancient individuals, with individuals being most closely related to *V*.*v*.*vicugna* or the llama/*L*.*g*.*cacsilensis* clade. Only individual C72 was found to be within the llama/*L*.*g*.*cacsilensis* clade.

### Ancestry proportions

To obtain high-resolution ancestry profiles, clarify the origins of each specimen, and assess possible interspecific gene flow, we calculated ancestry proportions for eight ancient individuals using admixfrog ^27^. Investigations using simulated aDNA damaged modern individuals showed a good ability to discern subspecies designation in the wild individuals using this approach as the highest proportion of single subspecies ancestry in each individual was as expected (Supplementary Fig S4). In the llama we saw a relatively high proportion of ancestral camelid DNA (0.11 vs. 0.01 in the wild species) but negligible levels of *Vicugna* ancestry (0.01). However, in the alpaca, we saw a much higher proportion of ancestral camelid DNA (0.44) as well as relatively high levels of *Lama* ancestry (0.13).

The empirical ancestry proportion results were in line with the PCA and phylogenetic tree results (Fig 2C and D). C72 had more *L*.*g*.*cacsilensis* ancestry relative to *L*.*g*.*guanicoe* compared to the other ancient *Lama sp*. individuals (Fig 2C). When looking into *Lama* sp. specific ancestry, C72 contained relatively higher levels of shared ancestry with modern llama than the other three investigated individuals (Fig 2D). Overall, the genetically identified *Lama sp*. individuals had more *L*.*g*.*cacsilensis* ancestry than *L*.*g*.*guanicoe* and *Vicugna sp* (Fig 2C). Putative *Vicugna* individuals have more *V*.*v*.*vicugna* ancestry than *V*.*v*.*mensalis* (Fig 2C). The small sized C69 contained mostly *L*.*guanicoe* ancestry, and the big size C38 contained mostly *V*.*vicugna* ancestry. Only a very small proportion of each individual contained ancestral South American camelid ancestry <0.04 (Fig 2C).

When using the wild species reference panel, the vast majority of the oldest individual’s, C46 from Tulán-52, ancestry could not be determined beyond ancestral *Lama* with the highest proportion of single subspecies ancestry (2%) coming from *L*.*g*.*cacsilensis*, the next closest being only 0.1% from *L*.*g*.*guanicoe*. When using the *Lama* only reference panel we found 48.9% of its ancestry as ancestral *Lama*, 50.1% as ancestral *L*.*g*.*cacsilensis*/llama, 0.58% as *L*.*g*.*cacsilensis* only, 0.25% as llama only, and 0.13% as *L*.*g*.*guanicoe* only.

### D-statistics

To further test the taxonomic placement of the archaeological specimens and to identify any signatures of interspecific gene flow we used D-statistics. However, very low-coverage ancient DNA data has been shown to introduce biases in D-statistics analyses, especially when mapping to a divergent reference genome ^28^. To address potential biases in our data, we conducted a simulated dataset analysis. Placing an individual with simulated ancient DNA damage into the H3 position produced comparable results to when placing a high quality version of the same individual in the H3 position (Supplementary Figure S5). Therefore, we could proceed with confidence knowing ancient DNA damage would not bias our topology/population structure results.

D-statistics results using the topology H1=*Lama sp*. H2=*Vicugna sp*. H3=Ancient individuals returned results consistent with the PCA, phylogenetic tree, and ancestry proportion analysis; individuals C75, C72, C89, C21 were more closely related to *Lama sp*. and individuals C50, C39, C38, and C20 were more closely related to *Vicugna sp*. (Fig 3A). Further investigations into the *Lama sp*. individuals showed that most had an overall closer relation to *L*.*g*.*cacsilensis* relative to llama or *L*.*g*.*guanicoe* (Fig 3B). However, C72 had similar levels of *L*.*g*.*cacsilensis and L*.*glama* ancestry, as seen by few significant results when calculating D-scores using *L*.*g*.*cacsilensis and L*.*glama* in the H1 and H2 position respectively. Further investigations into the *Vicugna sp*. individuals showed an overall closer relation to *V*.*v*.*vicugna* relative to alpaca or *V*.*v*.*mensalis* (Fig 3C).

**Figure 3:**
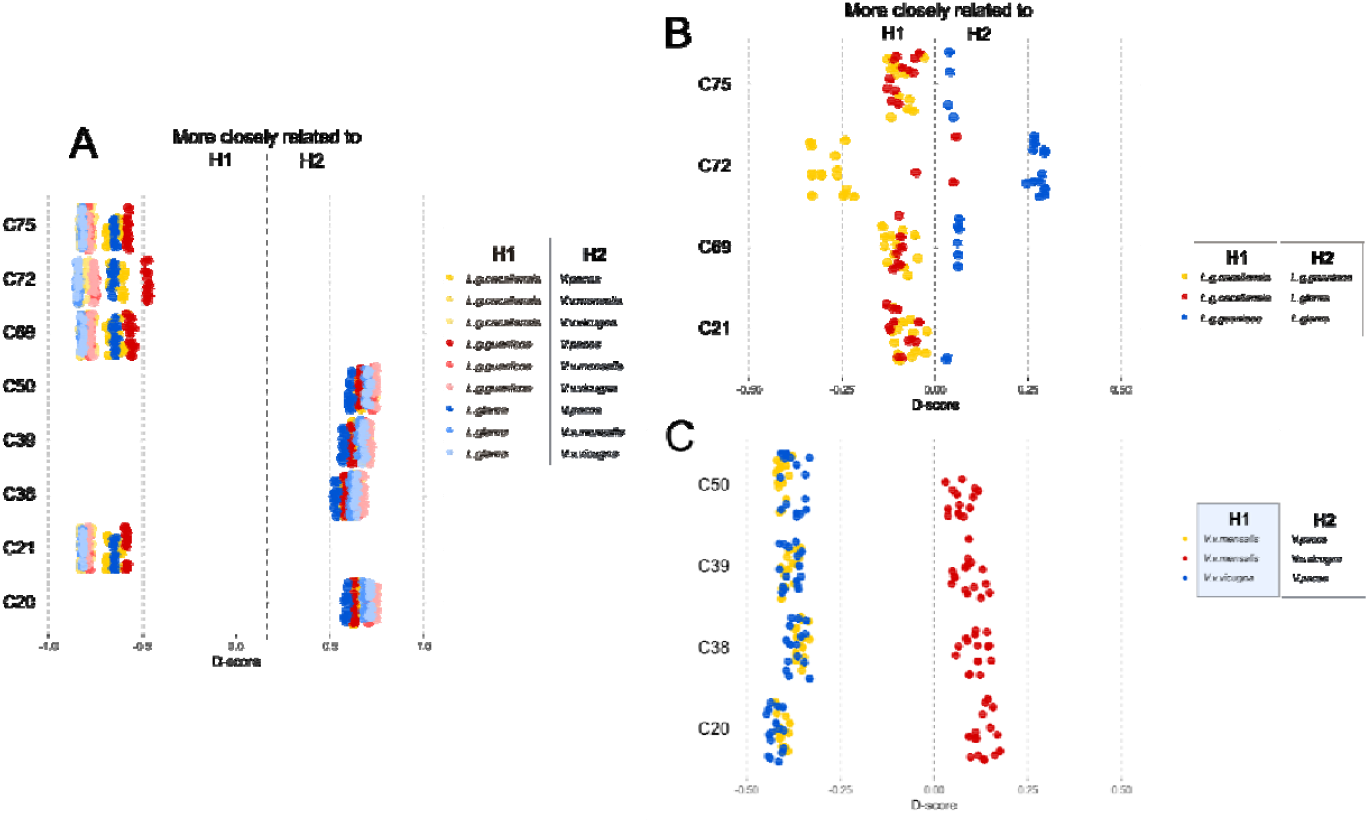
Test for closer evolutionary relationships between our ancient samples and the modern (sub)species using D-statistics and various input topologies. A) Comparisons between *Lama* and *Vicugna* B) Comparisons within *Lama* C) Comparisons within *Vicugna*. Non-significant values |Z|<3 not shown.

Comparing results placing a llama individual with simulated ancient DNA damage into the H2 position, a *Lama* sp. in the H1 position, and a *Vicugna* sp. individual in the H3 position to those when placing a high quality version of the same llama individual in the H2 position, we saw a slight discrepancy in the D-scores. The mean deviation was 0.0322 which we used to correct for ancient DNA related biases in the interpretations of the empirical data (Supplementary Figure S6).

Investigations into the topology ((*Lama* sp., C21), *Vicugna* sp.) found that individual C21 had less gene flow with *Vicugna* sp. than all modern llama individuals, similar levels of gene flow with *V*.*v*.*vicugna* and *V*.*v*.*menalisis* but more with alpaca compared to *L*.*g*.*guanicoe*, and less gene flow with alpaca but similar levels with *V*.*v*.*vicugna* and *V*.*v*.*menalisis* compared to *L*.*g*.*cacsilensis* (Fig 4). Investigations into the topologies ((*Lama* sp., C72), *Vicugna* sp.) found that Tulan individual C72 had less gene flow with *Vicugna* sp. than modern llama, similar levels of gene flow with *V*.*v*.*vicugna* and *V*.*v*.*menalisis* but more with alpaca compared to *L*.*g*.*guanicoe*, and comparable levels with all *Vicugna* sp. compared to *L*.*g*.*cacsilensis* (Fig 4).

**Figure 4:**
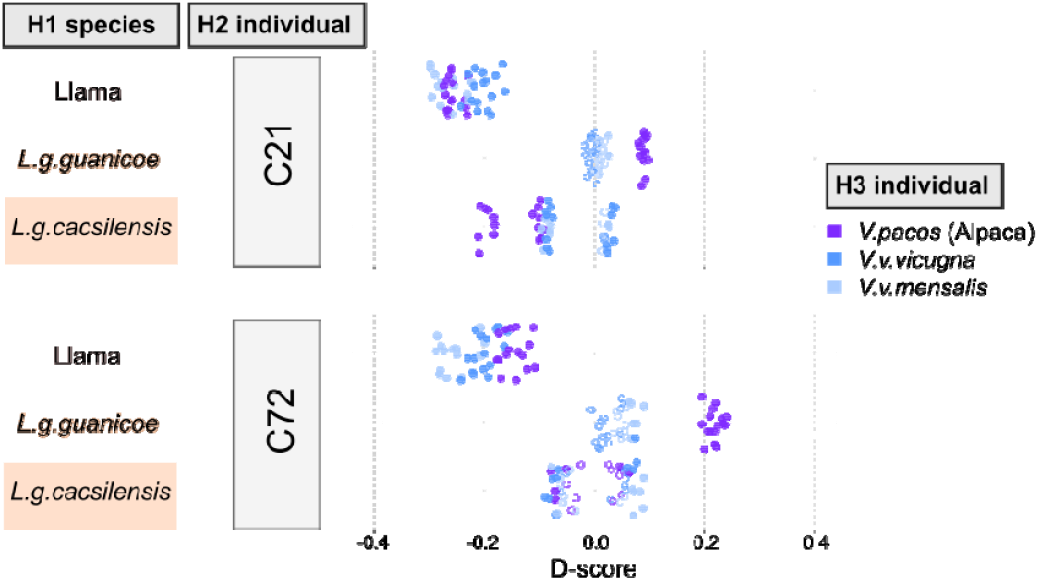
Test for gene flow between ancient individuals and Vicugna sp. using D-statistics. A negative D-score shows relatively more gene flow between H1 and H3 compared to H2 and H3, whereas positive shows relatively more gene flow between H2 and H3. Non-significant values |Z|<3 are shown with open circles.

### Genetic sex determination

To evaluate whether the sex distributions in our archaeological assemblages reflect hunting or herding strategies, we genetically sexed all individuals with >5,000 mapped reads using the SeXY pipeline ^29^. We were able to genetically determine the sex of 32 males and 29 females (Supplementary table S2). We were unable to determine the sex of 14 individuals due to either low number of mapping reads or ambiguous results. Further separating into each site, we found: Tulán-52 (1 F), Tulán-54 (24 F and 26 M), Tulán-85 (6 M and 3 F), and Tulán-94 (1 F). Further splitting by taxonomic status, we found similar proportions of male and female *Lama* at Tulán-54, slightly more females than male *Vicugna* at Tulán-54, and more males than females at Tulán-85 for both genera (Table 1). The one individual that showed the highest proportion of ancestry shared with modern llama (CL72) was a male. We note that while our findings provide insights into sex ratios at these sites, the limited sample size, especially at Tulán-85, presents a challenge in making broad inferences.

**Table 1:**
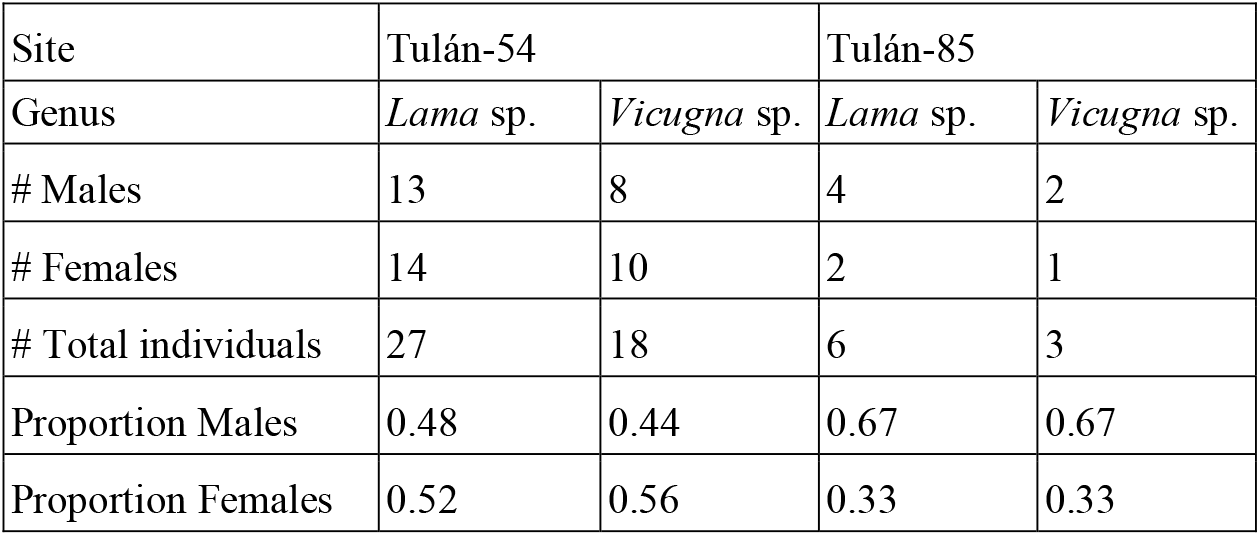
Genetic sexes of the individuals with known taxonomic status. Taxonomic status was based on the palaeogenomic results unless <0.001x genome-wide coverage then it was based on the osteological measurements - small = *Vicugna*, big = *Lama*.

## Discussion

The Tulán ravine, occupied during the Late Archaic and Early Formative periods, contains both ceremonial and residential sites with archaeological evidence for camelid exploitation, fibre processing, and ritual activity. Investigating palaeogenomic data, we identify the genetic ancestry and sex of camelids from these contexts and interpret these findings in light of the archaeological record to better understand the composition of camelid populations present during this critical period of economic and social transition in the South-Central Andes. Specifically, we investigated 49 individuals with mean genome-wide coverages >0.001x from the ceremonial sites of Tulán-52 (Late Archaic; n=1) and Tulán-54 (Early Formative; n=40) as well as the residential site of Tulán-85 (Early Formative; n=8). We found evidence for both genera, Vicugna and Lama, in our dataset. Further filtering to 26 individuals >0.01x showed that all individuals, except one, likely represented ancestry not found in modern individuals (Fig 2). Moreover, while our ancient data quality undoubtedly presented some limitations, particularly due to the low coverage and poor DNA preservation typical of temperate regions ^30^, careful evaluation of biases through simulated data demonstrated that even low-coverage palaeogenomes can yield valuable insights when these biases are accounted for. This highlights the potential of low quality data to provide meaningful results, even from challenging archaeological contexts.

The *Vicugna* individuals we identified most likely belonged to the subspecies *V*.*v*.*vicugna*, which was not unexpected given that the local subspecies of *Vicugna* is *V*.*v*.*vicugna* ^31^. The presence of small individuals with characteristics indicative of vicuñas in the Tulán sites may reflect the value of vicuña fibre, which has been appreciated since pre-Hispanic times ^7^, and is still greatly appreciated today. This may be the reason that most of the small animals in Tulán correspond to vicuñas. The use of camelid fibre is supported by microscopic camelid fibre analyses^32^. While the overwhelming majority of fibres were light brown natural colours, a small quantity of pure black fleece suggests colour variation, hinting at early colour selection of pre-domesticated camelids ^33^. While this could suggest a lost domesticate-like lineage filling a similar role as the alpaca, this could also be the result of dye so should be considered with caution ^34^.

*Lama* individuals were more difficult to place than the *Vicugna*. All individuals had a tendency towards *L*.*g*.*cacsilensis* but all bar one had considerable levels of *L*.*g*.*guanicoe* ancestry. The current distribution indicates that *L*.*g*.*cascilensis* are generally found further north but may slightly overlap with *L*.*g*.*guanicoe* in Northern Chile ^35,36^. Therefore, the ancient Tulán individuals may represent a locally extinct *L*.*g*.*cacsilensis* population that hybridised with the local *L*.*g*.*guanicoe* individuals. The one individual more closely related to *L*.*g. cacsilensis*, with relatively high levels of llama ancestry, is particularly interesting from a domestication standpoint. When placed in the broader archaeological context of the Tulán ravine, where camelids appear to have been used in a domesticated role, these genetic findings support the idea that this individual may represent either an early domesticated llama or a wild population ancestral to contemporary llamas, with full domestication likely occurring later. Being an early domesticated llama may have also been the case for our one sample from the Late Archaic Tulán-52 individual (C46) which contained mostly *L*.*g*.*cacsilensis/*llama ancestry with relatively little presence of *L*.*g*.*guanicoe*. There is a cultural continuity between the Late Archaic and the Early Formative, so it is to be expected that these populations continued to develop the domestication/specialisation process. However, the very low coverage (0.008x) nature of this sample prevented us from delving deeper. Taken together, our results suggest individuals that were not the direct ancestors of contemporary domesticated species were the main source of South American camelids used in both domestic and ceremonial activities in the Tulán ravine during the Early Formative period (3,360–2,370 cal. yr BP).

Osteometric analyses of the excavated samples of Tulán-54 previously revealed the presence of the two wild species: guanaco and vicuña, and the domestic species, llama ^26^. However, the large degree of overlap between domestic and wild individuals makes clear results difficult to obtain ^37^. Taking these difficulties together with the fact that there are other lines of evidence (e.g. artefacts and fibres ^38,39^) supporting the notion that there were domesticated animals in the Tulan ravine since the Late Archaic period ^40^, suggest the large animals were utilised in a domesticated role despite sharing different ancestry to contemporary llama. As it has been hypothesised that llama domestication was a complex process and may have occurred in several different areas of the Andes ^16^, the Tulán individuals could represent a domestic lineage that was subsequently replaced or diluted through admixture with other populations. Similar patterns of replacement through introgression have been documented in other domestic mammals, such as pigs, where Near Eastern domestic ancestry was largely replaced by European wild boar ancestry following introduction ^41^, and horses, where ancient domestic lineages were replaced by later lineages of different origin ^42^. While the exact mechanisms and timing may differ, these cases illustrate how domesticated lineages can be transformed or supplanted through subsequent gene flow.

A previous study using mitochondrial genomes of the same individuals suggested that individuals whose size did not match their maternal ancestry (e.g. small size but *Lama* ancestry or big size but *Vicugna* ancestry) may have been hybrid individuals, which was interpreted as indicating domesticated individuals ^17^. Our palaeogenomic analyses find that these potential hybrid individuals do not contain clearly higher levels of shared ancestry between genera. This finding challenges the assumption that mitochondrial and nuclear signatures, as well as osteometric measurements, should align to accurately determine domestication status. As mitochondrial DNA reflects only the maternal lineage, it may not fully capture genetic complexities such as intergenerational hybridisation and introgression. Moreover, osteometric measurements alone, though valuable in identifying size variation, do not account for the possibility of mislabelling or misidentification of juveniles versus adults. As such, we propose that the apparent discrepancy between mitochondrial and nuclear data regarding hybrid status is likely due the inherent limitations of mitochondrial analysis. Consequently, the assumption that size, mitochondrial ancestry, and nuclear signatures should consistently align may not always hold due to the complexity of domestication processes and the potential for interbreeding between different camelid species and subspecies over time. While mitochondrial introgression may have occurred, a single sporadic mitochondrial introgression event could lead to inferences of widespread hybridisation, which, as shown by our nuclear genomic analysis, is not supported.

Despite early Spanish observers acknowledging the importance of camelid pastoralism, they generally failed to distinguish between domestic species in their records. This contrasts with *Lama* and *Vicugna* having long been distinct, biologically divergent genera separated approximately 2–3 million years ago ^20^ and culturally differentiated by pre-Hispanic communities ^4^. The subsequent indiscriminate hybridisation between domesticates resulted in a loss of their exquisite fibre qualities and colourations that made them so prized in prehistoric Andean communities ^4^. This uncontrolled hybridisation between domesticated species and the population decline post-conquest makes identification of pre-hispanic genetic ancestry of modern individuals difficult. Previous work exclusively analysing genomes from modern individuals has found both llama and alpaca to contain evidence of intergeneric gene flow tracing back to ∼500 years ago ^20^. Our results show that the putative llama in our dataset contained less *Vicugna* ancestry than modern llama, and similar amounts to wild *L*.*g*.*cacsilensis*, the likely ancestral (sub)species of modern llama. Therefore, we suggest that intermixing between *Vicugna* (most probably alpaca) and llama was likely not a key component of the early domestication process, and only occurred within the last ∼3,000 years.

While the absence of evidence for early hybridization in our samples does not exclude its sporadic occurrence, it suggests that hybridisation was not a widespread practice during the studied period. Palaeogenomic data from more domesticated individuals will undoubtedly help shed further light on this issue.

In addition to the insights gained from the ancestry analysis, we were also able to infer the sex of the individuals using the palaeogenomic data. South American camelids are sexually monomorphic ^43^ showing sexual dimorphism when specific bones are available (e.g. pelvis and canine teeth) ^44^. While it is possible to sex South American camelids, this can rarely be done due to the fragmentary nature of most assemblages. Therefore, the palaeogenomic data presented a unique opportunity to make inferences on sex. Phalanxes and Astragali have been used in this study, due to their high frequency, completeness, and in the case of the former, as their fusion stage indicates the presence of adult animals (over 24 months) ^45^. Thus, our results examine only the adult sex ratio in archaeological assemblages.

Sex distribution in archaeological assemblages reflects both hunting and herding strategies. Hunting assemblages are generally expected to be dominated by adult remains, with sex ratios balanced or male-biased, whereas herding assemblages often include an adult sex ratio skewed toward females ^46^. Herding practices can vary, for example, through selective culling of infertile or undesirable males, which may produce more balanced adult sex ratios ^47^. In the Tulán assemblages, we observe roughly similar proportions of males and females across sites and genera (Table 1), although the small sample from Tulán-85 requires cautious interpretation. These patterns are consistent with either hunting or herding with selective male culling and align with our genomic findings showing that most individuals at these sites were wild individuals and not the direct ancestors of domesticated camelids.

## Materials and Methods

### Archaeological contexts

It is proposed that the location of Archaic and Formative settlements in the Tulán area reflects a combination of favourable cultural, social, and environmental conditions. These conditions supported a shift from a society primarily reliant on hunting and foraging, characterized by seasonal mobility, small group sizes, and exploitation of wild resources, to one increasingly oriented toward pastoralism, marked by the management of domesticated camelids, more permanent settlement structures, and reliance on animal products for subsistence.

The Late Archaic period (ca. 5290-4090 cal. yBP) is marked by large campsites like Tulán-52, the emergence of substantial architecture with clusters of circular structures, and a diversified, innovative lithic industry. This period also saw the appearance of rock art, long-distance interactions, and a subsistence strategy that combined hunting with early camelid domestication. Altogether, these features point to a growing sociocultural complexity ^48^.

The Early Formative period (ca. 3360 - 2370 cal. yBP) is characterised by the emergence of large settlements with a complex and planned ritual architecture along with the intensification and expansion of productive practices, an emphasis on ritual expressions, the appearance of new technologies such as pottery, and three large contemporary Formative settlements (Tu-54, Tu-122 and Tu-85) have been documented for this period. These are located within a 15 km area between the border of the Salar de Atacama and the Tulán ravine, at an altitude ranging from 2300–3200m. These sites have different functions. At Tulán-122, located in the middle sector of the ravine, residential aspects were emphasised. In contrast, Tulán-85, is located on the border of the resource-rich salt flat. These settlements shared a common political and religious organisation centred around the Tulán 54 site, the only site that possesses monumental architecture ^25^.

The sunken temple structure identified in Tulán-54 at the centre of a refuse mound contained 27 human infant graves accompanied by offerings, rock art, structured fireplaces, and offering pits, among other elements, suggesting that the structure had a ceremonial function. The main structure identified at the site is a large, sunken semi-oval ceremonial structure surrounded by a perimeter wall made of large, vertically positioned stone blocks. The inner space is divided into six precincts, with a single oval structure in the middle ^25,26^. The recovered camelid anatomical units display a very similar temporal and spatial pattern across all occupations and among the different precincts of the ceremonial structure, which was filled intentionally with the remains of butchering, processing, consumption and feasting activities. The waste was likely generated outside and then ritually incorporated ^49^. On the border of the Salar de Atacama, Tulán 85 corresponds to an extensive occupation characterised by dense domestic monticulated deposits. The site also includes marginal structures and a sector with burials of newborns ^50^. Finally, Tulan-94 site is composed of groups of circular and sub-circular structures, with a total of 19 living enclosures. The evidence suggests the transitional character of the site, since it has material components corresponding to the Late Archaic and Early Formative phases ^25^.

### Samples

All research was conducted in accordance with Chilean regulations. Excavation permits were issued by the Consejo de Monumentos Nacionales de Chile (CMN) under permit Nº 4409 (18.11.2013). Export of osteological samples (Camelidae) to the University of Copenhagen for ancient DNA analysis was authorized by CMN ordinances ORD. Nº 03642 (21.10.2016), Nº 03746 (27.10.2016), and Nº 03425 (20.01.2017), within the scope of FONDECYT project 1130917. All exported samples were photographed, labeled, and transported via certified courier in accordance with CMN requirements. Although none of the camelid remains analysed here were directly radiocarbon dated, chronological placement is based on radiocarbon dates from associated archaeological contexts (e.g., stratified deposits, fireplaces, and burials) ^51^.

Our study consisted of 75 ancient individuals sourced from four archaeological sites: Tulán-52, Tulán-54, Tulán-85, and Tulán-94, and have previously undergone mitochondrial and morphological analysis ^17^ (Supplementary tables S1 and S2). In brief, samples were collected from both ceremonial and domestic contexts across different sectors of the archaeological sites to capture spatial and temporal variation. Bone elements (primarily first anterior phalanges and astragali) were selected from stratified excavation units and structures, with care taken to sample different individuals. Only adult specimens, identified by fused epiphyses or advanced ossification, were included in the morphometric analysis. Measurements were taken using Vernier calipers to the nearest 0.1 mm, following established osteometric protocols, and were recorded prior to aDNA sampling to avoid damage-related bias. Key dimensions were measured on each bone type to assess size variation, and results were compared against published morphometric data for all four modern South American camelid species to evaluate patterns of morphological differentiation. The threshold for distinguishing small and large individuals was determined based on a comparison of key osteometric measurements from the first anterior phalanx (breadth of proximal articulation [BFp] and depth of the proximal epiphysis [Dp]) and the astragalus (breadth distal [Bd] and greatest length medial [GLm]). The distinction between small and large groups was tested statistically by correlating measurements from both the phalanx and astragalus using PAST ^52^. Published morphometric data for all four modern South American camelid species were used to establish the thresholds for size differentiation ^53,54^.

To build a genomic reference panel, we also included published genomic data from 80 modern individuals (DNAzoo.org, ^20,35^). The modern individuals represented all extant *Lama* and *Vicugna* subspecies (Figure 1).

### Bioinformatic processing

The raw sequencing data for the ancient Tulán individuals was generated in a previous study where only the mitochondrial DNA was analysed and obtained from the authors ^17^ while the raw sequencing reads for the modern individuals were downloaded from the European Nucleotide Archive (ENA) (Supplementary tables S1 and S2). For all individuals, we removed Illumina adapter sequences, low-quality reads (mean q <25), short reads (<30bp), and merged overlapping read pairs with Fastp v0.23.2 ^55^. We mapped the processed reads (merged only for the ancient individuals) to the alpaca reference genome (Genbank accession: GCF_000164845.3) using Burrows-wheeler-aligner (BWA) v0.7.15 ^56^ and either utilising the aln algorithm, with the seed disabled (-l 999) (otherwise default parameters) for the ancient individuals or the mem algorithm with default parameters for the modern individuals. We chose the alpaca genome as the reference genome due to its hybrid ancestry ^20^, making it putatively more suitable when making genomic comparisons between genera. We parsed the alignment files and removed duplicates and reads of mapping quality score <30 using SAMtools v1.6 ^57^. We checked for ancient DNA damage patterns of the ancient individuals using mapdamage v2 ^58^. As the ancient individuals had also undergone targeted enrichment for mitogenomes and some selected genes, we aligned the probes to the reference genome using BLAST v 2.15.0 ^59^ using default parameters and we removed any regions of the alpaca genome aligning to the probes from the mapped bam files using bedtools v2.29.1 ^60^.

### Genetic sex determination

We identified scaffolds putatively originating from the sex chromosomes in the alpaca assembly by aligning the assembly to the alpaca X (included in the reference genome) and Human Y (Genbank accession: NC_000024.10) chromosomes using satsuma synteny v2.1 ^61^ with default parameters. We followed the SeXY pipeline ^29^ to identify the sex of each of the ancient individuals. We set a minimum threshold of 5,000 mapped reads to ensure the reliability of the results. If an individual had a mean X:A ratio of <0.7 it was designated as a male. If an individual had a mean X:A ratio of >0.8 it was designated a female. Individuals with mean ratios between 0.7 and 0.8 were deemed undetermined.

### Ancient DNA Simulation

To assess the impact of ancient DNA damage on our analyses we simulated the damage patterns and read lengths of individual C21 onto representatives of each modern subspecies (seven in total) using gargammel v1.1.4 ^62^. We chose C21 due to it having the highest coverage of the ancient samples. To do this, we generated consensus fasta files from the mapped bam files of the modern individuals using ANGSDv0.921^63^ and the following parameters -dofasta 2, minimum read depth of 5 (-setmindepthind 5), and minimum mapping and base qualities of 30 (-minq 30 - minmapq 30). The modern individuals included *L*.*glama* (Llama2 and Llama3), *V*.*pacos* (Alpaca3), *L*.*g*.*cacsilensis* (Cacsilensis3), *L*.*g*.*guanicoe* (Guanicoe3), *V*.*v*.*mensalis* (Mensalis2), and *V*.*v*.*vicugna* (Vicugna1). We mapped the simulated ancient DNA reads to the alpaca reference genome following the same approach as for the ancient Tulán individuals. We additionally downsampled them to ∼0.01x using SAMtools.

### Principal Component Analysis

We evaluated the relationships between the modern and ancient camelids included in this study by performing PCAs using a pseudohaploid base call with ANGSD. We initially did this for two different sample sets, either for all modern and ancient individuals >0.01x or all modern and ancient individuals >0.001x. We computed consensus pseudohaploid base calls specifying the parameters: minimum mapping and base qualities of 20 and 30 respectively (-minmapQ 20 -minQ 30), calculate genotype likelihoods using the GATK algorithm (-GL 2), calculate major and minor alleles based on genotype likelihoods (-doMajorMinor 1), remove transitions (-rmtrans 1), only consider autosomal chromosomes > 10Mb (-rf), skip triallelic sites (-skiptriallelic 1), only consider reads mapping to one region uniquely (-uniqueonly 1), make a consensus identity by state base calls (-doIBS 2), output a covariance matrix (-doCov 1) and only consider variable positions where the minor allele occurs in at least two individuals (-minminor 2). We also repeated this analysis but with only individuals belonging to the *Lama* genus. We set the minimum individual threshold to the number of modern individuals +1, i.e. 81 for the complete dataset and 57 for the *Lama* only.

### Phylogenetic Analysis

To build a neighbour joining tree, we first constructed a distance matrix, which we output from ANGSD when performing PCA (setting the parameter -makematrix 1). We converted the distance matrix into a neighbour joining phylogenetic tree using fastME v2.1.6.1 ^64^ and default parameters.

### Ancestry proportions

We calculated the ancestry proportions of eight ancient individuals using admixfrog v0.7.2^27^. We included four big size camelids (C21, C38, C72, C75) and four small size camelids C20, C39, C50, C69. We selected these based on their higher coverage or unexpected placement within the PCA. Individuals C69 and C38 did not correspond to what was expected based on their size and C75 clustered differently compared to the other big sized ancient individuals. We also ran this analysis on the lone individual from Tulán-52 (C46) as it comes from an earlier time period which may be closer to the onset of domestication. Initially we only considered the wild species as the ancestry states, each represented by a single individual; *L*.*g*.*cacsilensis* - Cacsilensis4, *L*.*g*.*guanicoe* - Guanicoe1, *V*.*v*.*mensalis* - Mensalis4, and *V*.*v*.*vicugna* - Vicugna2. We repeated the analysis for the ancient *Lama* individuals using the same two *Lama sp*. subspecies as above plus Llama9 as ancestry states. As input for the reference panel we created a multi-individual variant call file with using BCFtools v1.15 ^65^, specifying autosomal scaffolds >10Mb (-f) and minimum mapping and base qualities of 20 (-q 20 -Q 20). We ran admixfrog specifying a minimum read length of 30bp, a bin size of 50kb, and otherwise default parameters. We only considered a bin in interpretations if it contained more than five SNPs. We pooled regions showing mixed ancestry between *Lama sp*. and *Vicugna sp*. as ancestral camelid in the first run and pooled regions showing any mixed ancestry in the *Lama sp*. only run as ancestral. Furthermore, to assess the accuracy and suitability of this method on our dataset, we also performed the same analysis (all wild (sub)species reference panel) on the modern individuals with simulated ancient DNA damage.

### D-statistics

D-statistics calculates shared derived alleles between non-sister branches of a given topology [[[H1, H2], H3], Outgroup]. An allele is considered as derived if it is different to the outgroup allele. Although it is most commonly implemented to investigate differential levels of gene flow between non-sister taxon, it can also be used to investigate shared ancestral polymorphisms and therefore topology/population structure ^66,67^.

We subsampled the dataset to include four modern individuals per subspecies and only the eight ancient individuals also used in the ancestry proportions analysis above. We calculated the D-statistics using a random base call approach in ANGSD (-doabbababa 1), specifying only autosomal scaffolds >10Mb and the following parameters: -minmapQ 30 -minQ 30 -blocksize 1000000 - rmtrans 1 -uniqueonly 1. To infer the ancestral sequence (-anc) we used a wild bactrian camel (*Camelus ferus*; NCBI biosample SRR1947250) which we also mapped to the alpaca genome following the approach mentioned above. We summarised the results using a block jackknife approach with the Rscript available in the ANGSD toolsuite (ANGSD_jackknife.R).

We filtered the output into three different sets. First to investigate which genus each ancient individual was most closely related to, we filtered for H1=*Lama sp*. H2=*Vicugna sp*. H3=Ancient individuals. Based on these results we then looked into which subspecies the ancient individuals were most closely related to with either H1=*Lama sp*. H2=*Lama sp*. H3=Ancient *Lama sp*. individuals or H1=*Vicugna sp*. H2=*Vicugna sp*. H3=Ancient *Vicugna sp*. Individuals. To investigate gene flow between the ancient *Lama sp* and *Vicugna sp*., we extracted results corresponding to the topologies H1=*Lama* sp., H2=C21, H3=*Vicugna* sp. H1=*Lama* sp., H2=C72, H3=*Vicugna* sp.

To investigate whether the ancient DNA damage could cause biases in the topology test results, we compared results when placing the individuals with simulated ancient DNA damage in H3 with the results generated using the high quality version of the same individual. To evaluate the ability of our dataset to infer gene flow between the putative ancient llama and *Vicugna* species, we also performed D-statistics with the two 0.01x simulated aDNA llama (Llama 2 and 3) individuals in the H2 position, *Vicugna sp* in the H3 position and the remaining *Lama* sp. individuals in the H1 position. By comparing these results to those when the high quality versions of llama 2 and 3 were in the same position, we were able to calculate a correction factor to account for potential biases. We found a mean deviation between the two results of 0.0322 (Supplementary figure S6) which we subtracted from the D-score and using the standard error output with ANGSD, we recalculated the Z scores.

## Acknowledgements

We would like to thank Anders Hansen for support in generating the sequencing data and Alba Rey-Iglesia for help curating the data. We would also like to thank Lautaro Núñez who through numerous projects (FONDECYT 1020316, 1070040 and 1130917) recovered the analysed camelid bone remains.

## Data availability

Raw sequencing data is available under NCBI BioProject ID: PRJNA1215279. Coordinates of the capture probes in the alpaca reference genome can be downloaded from https://erda.ku.dk/archives/7d1d408b1711f5ab7e34a845bf58f9a8/published-archive.html

