## Supplementary information for "Palaeogenomics suggest domesticated camelid herding and wild camelid hunting in early pastoralist societies in the Atacama Desert"

**Supplementary Tables**

**Supplementary table S1:** Modern sample information and mapping statistics

**Supplementary table S2:** Ancient sample metadata, mapping statistics, SeXY values, and biometric data. BFp: Greatest breadth of the facies articularis proximalis; Bp: Greatest breadth of the proximal end; Dp: Greatest depth of the proximal end; DFp: Greatest depth of the facies articularis proximalis; A: Minimum depth of the epiphysis; GLm: Greatest length of the lateral half; Bd: Greatest breadth of the distal end.

**Supplementary table S3:** List of individuals used for the D-statistics analyses

**Supplementary Figures**

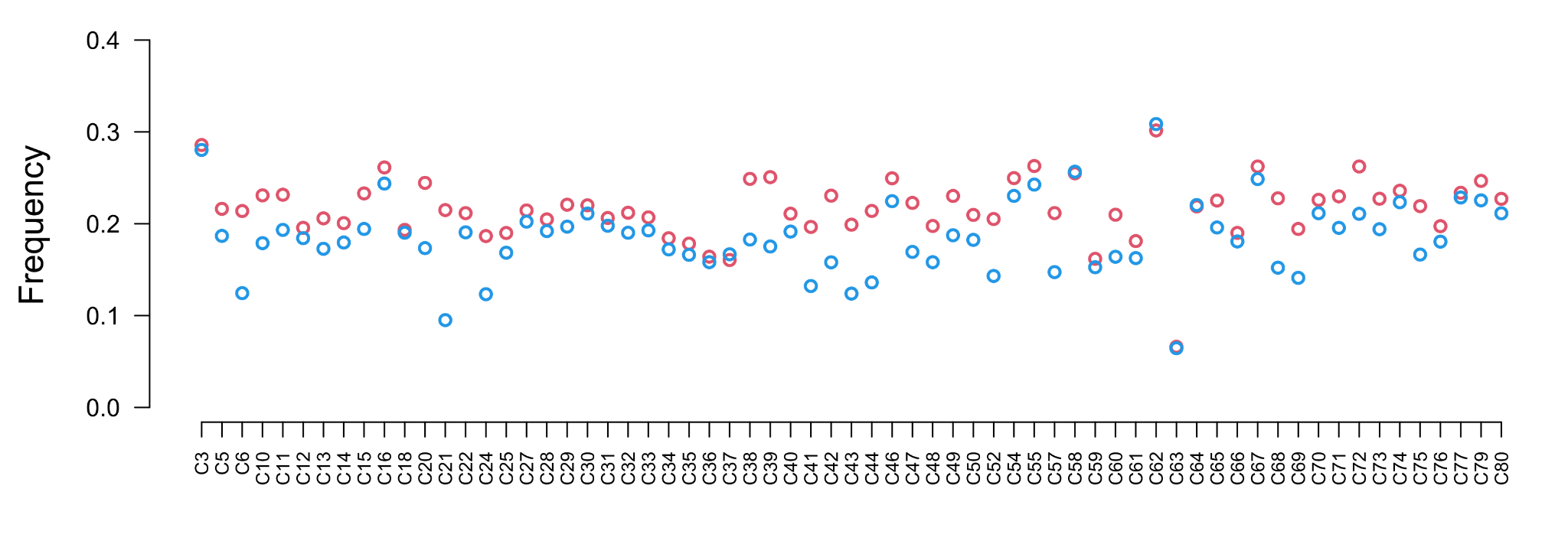

**Supplementary figure S1: DNA damage results taken from Mapdamage for all individuals with >5,000 mapped reads.** The frequency of C-T transitions on the first site from the 5-prime end of the read is shown in red, and G-A transitions on the first site from the 3-prime end of the read is shown in blue.

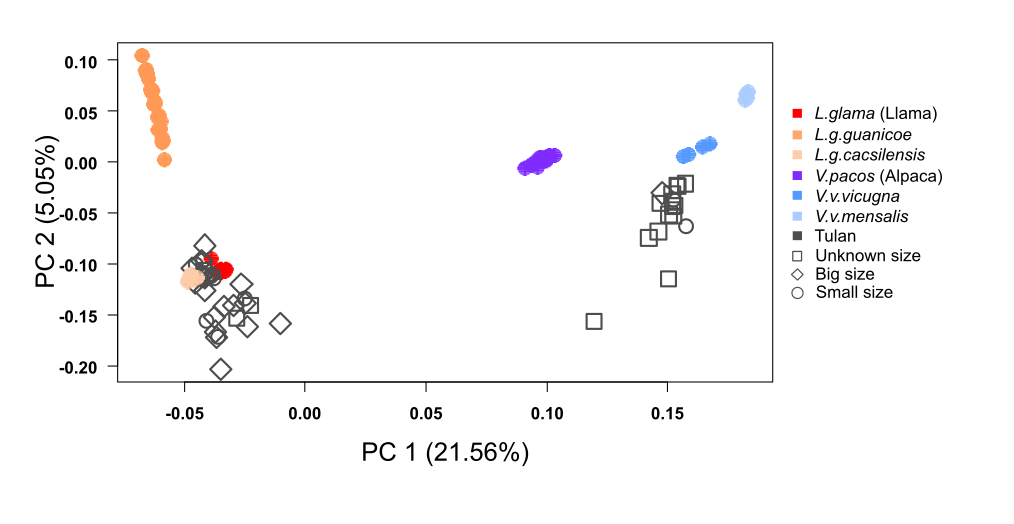

**Supplementary figure S2: Pseudohaploid PCA generated including Tulan samples >0.001x and the entire modern dataset.** Analysis performed using 5,122,839 transversion sites.

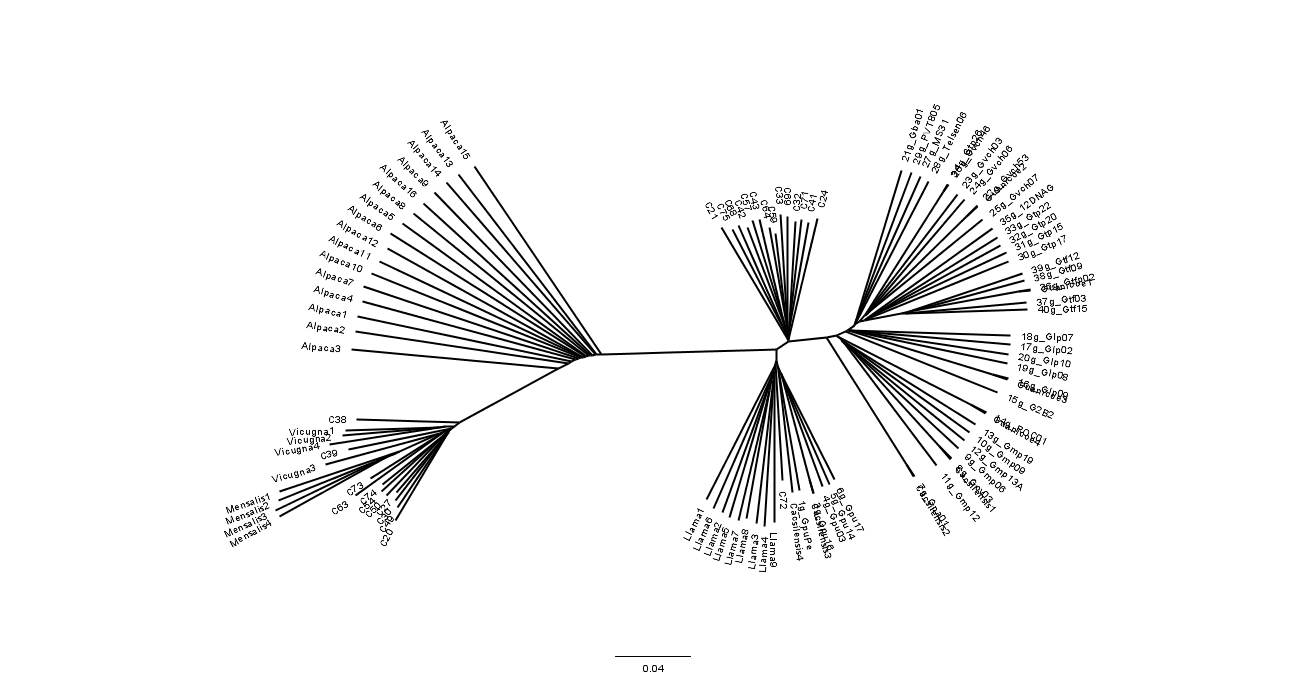

**Supplementary figure S3. Neighbour joining phylogenetic tree of all individuals >0.01x included in the study.** Scale bar indicates the proportion of identity by state differences.

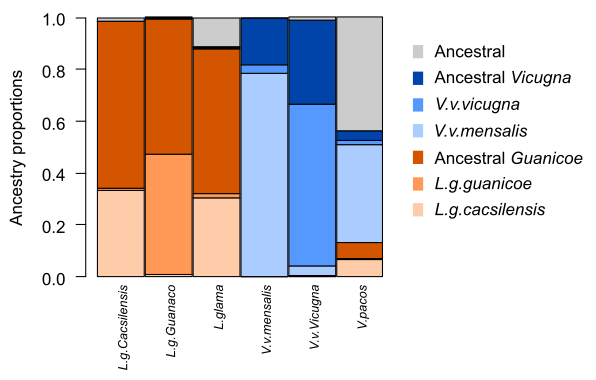

**Supplementary figure S4: Admixfrog results when using modern samples with simulated ancient DNA damage.** *L.g.cacsilensis* (Cacsilensis3), *L.g.guanicoe* (Guanicoe3), *L. glama* (Llama2), *V.v.mensalis* (Mensalis2), V.v.vicugna (Vicugna1), and *V. pacos* (Alpaca3). Ancestral indicates shared ancestry between *Lama* and *Vicugna*.

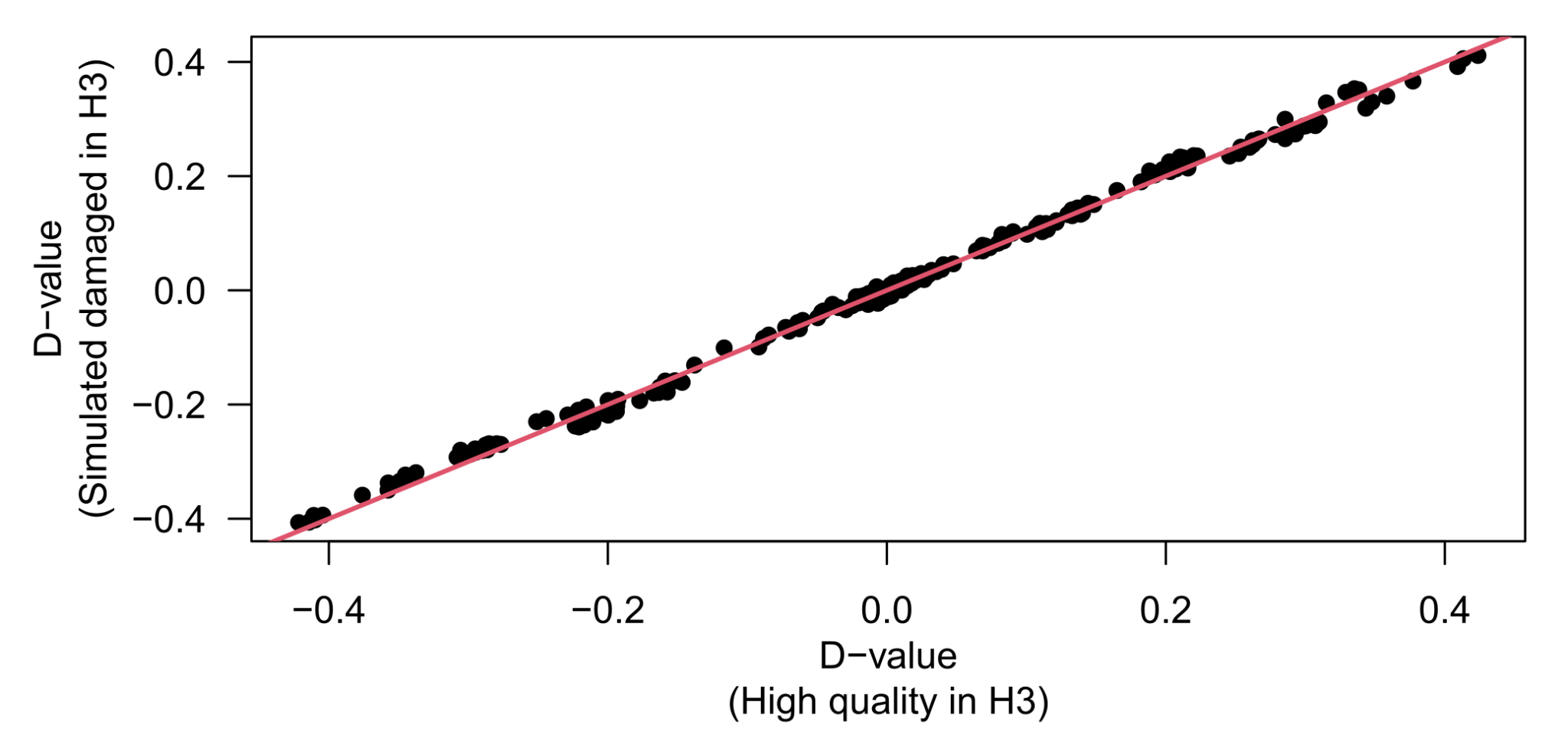

**Supplementary figure S5: Testing the robustness of D-statistics results when having a simulated ancient individual in the H3 position relative to the high quality version of the same individual.** Red line indicates a 1:1 result and therefore no bias.

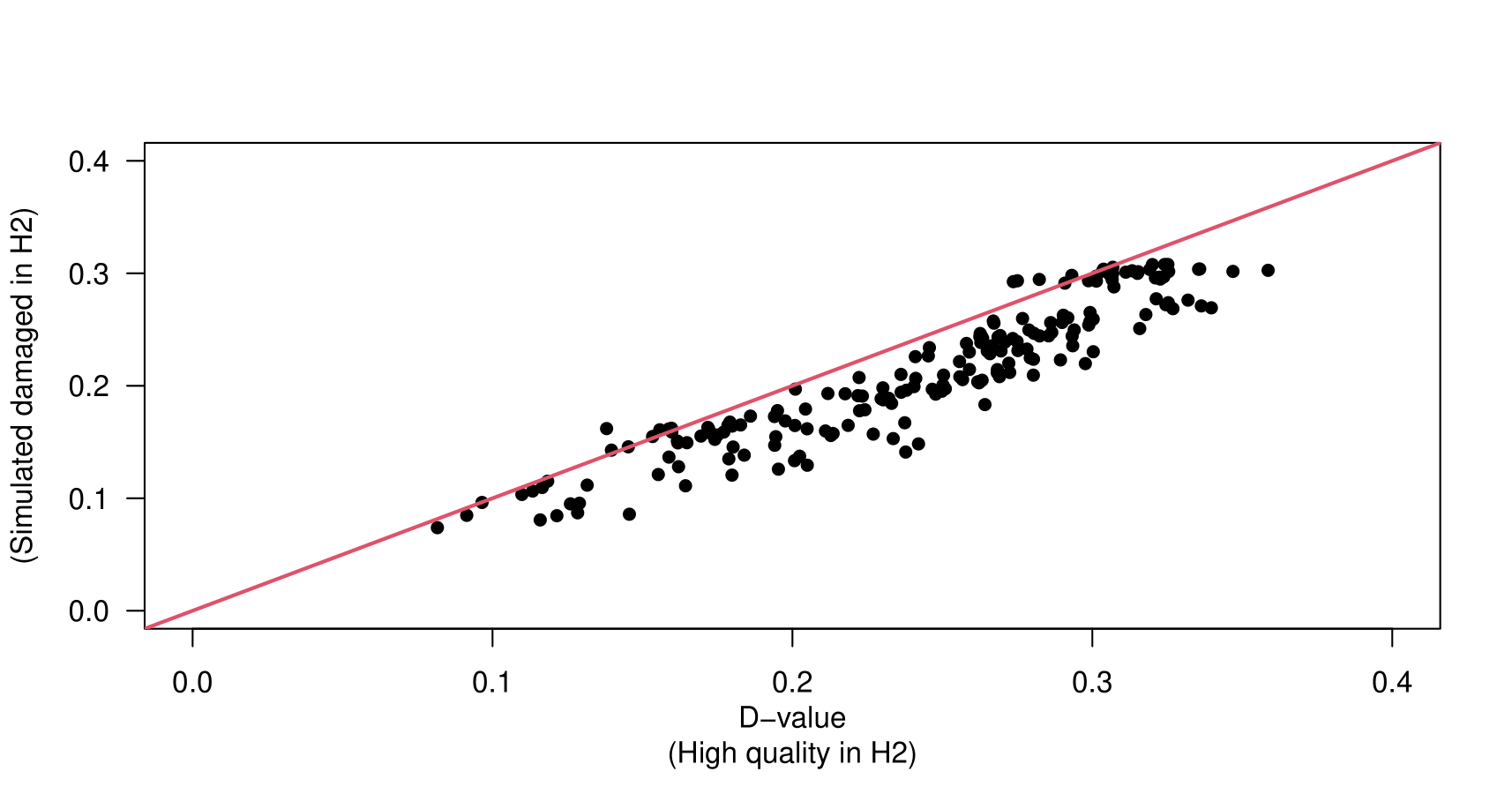

**Supplementary figure S6: Testing the robustness of D-statistics results when having a simulated ancient individual in the H2 position relative to the high quality version of the same individual.** This test only assessed results for the topology ((*Lama guanicoe/cacsilensis*, llama), Vicugna sp.). Red line indicates a 1:1 result and therefore no bias.
